# Creating outbred and inbred populations of haplodiploid mites to measure adaptive responses in the lab

**DOI:** 10.1101/2020.02.22.960682

**Authors:** Diogo P. Godinho, Miguel A. Cruz, Maud Charlery de la Masselière, Jéssica Teodoro-Paulo, Cátia Eira, Inês Fragata, Leonor R. Rodrigues, Flore Zélé, Sara Magalhães

**Affiliations:** Centre for Ecology, Evolution and Environmental Changes – cE3c, Faculdade de Ciências da Universidade de Lisboa, Edifício C2, 5º Piso, Sala 2.5.46. Campo Grande, 1749-016 Lisboa, Portugal

**Keywords:** experimental evolution, spider mites, genetic diversity, biological resources

## Abstract

Laboratory studies are often criticized for not being representative of processes occurring in natural populations. This can be partially mitigated by using lab populations that capture large amounts of variation. Additionally, many studies addressing adaptation of organisms to their environment are done with laboratory populations, using quantitative genetics or experimental evolution methodologies. Such studies rely on populations that are either highly outbred or inbred. However, the methodology underlying the generation of such biological resources are usually not explicitly documented.

Given their small size, short generation time, amenability to laboratory experimentation and knowledge of their ecological interactions, haplodiploid spider mites are becoming a widely used model organism. Here, we describe the creation of outbred populations of two species of spider mites, *Tetranychus urticae* and *T. evansi*, obtained by performing controlled crosses between individuals from field-collected populations. Subsequently, from the outbred population of *T. evansi*, we derived inbred lines, by performing several generations of sib-mating. These can be used to measure broad-sense heritability as well as correlations among traits. Finally, we outline an experimental evolution protocol that can be widely used in other systems. Sharing these biological resources with other laboratories and combining them with the available powerful genetic tools for *T. urticae* (and other species) will allow consistent and comparable studies that greatly contribute to our understanding of ecological and evolutionary processes.

## Introduction

The ultimate aim of evolutionary ecology should be to understand the processes that shape individual traits and ecological processes in natural populations. This can be achieved by studying populations in their natural environment. However, this approach suffers from the difficulty in controlling several environmental variables simultaneously. Laboratory studies, in contrast, while allowing for controlled variables, are often criticized for not being representative of the processes occurring in natural populations. Using laboratory populations that capture a large fraction of the natural variation found in field populations may be a form to mitigate this setback.

One of the most challenging tasks in evolutionary biology is to understand how populations adapt to particular biotic or abiotic stresses. Detecting such adaptations in the field is commonly done via reciprocal transfers or common garden experiments. These methods rely on comparing traits from individuals of different populations in their own vs the others’ environment, or in a common, controlled environment (Kawecki and Ebert 2004, Hereford 2009, Blanquart et al. 2013). However, these methods suffer from the shortcomings of comparative studies. Indeed, it is difficult to attribute differences in trait values to a specific environmental variable, as environments may differ in several dimensions. Additionally, disentangling the relative role of adaptation and correlated responses to selection in shaping traits is hampered by the fact that the ancestral state is often unknown (Lauder et al. 1993, Leroi et al. 1994). Although it is often assumed that the ancestral state is one of the extant forms (Lauder et al. 1993, Leroi et al. 1994), this assumption is often difficult to document or justify. By relying on contemporary data, erroneous conclusions may be drawn concerning the process of adaptation and its potential costs (Bell and Reboud 1997; Magalhães et al. 2009). These setbacks can be circumvented by sampling natural populations and introducing them to the laboratory, where they can be exposed to specific environmental variables, and their responses analysed using either quantitative genetics or experimental evolution.

Quantitative genetics uses several designs to evaluate the genetic vs environmental contribution to a particular phenotype. In all cases, however, it is important that the population used to infer these contributions is sufficiently variable. In particular, some designs rely on a panel of inbred lines, created from the same population. To ensure that this panel is composed of different genotypes, it is important to derive it from a highly outbred population. Such panel can then be used to measure the broad-sense heritability of a given trait, as well as genetic correlations between traits. Although the most famous and complete panels are found in Drosophila (DGRP – Mackay et al. 2012, DSPR - King et al. 2012), this resource has also been used in plants (Kover et al 2009, Wills et al. 2013) and other animals (Table 1).

**Table 1.** Inbred line panels in different animal species

| **Organism** | **Characteristics** | **# of lines** | **Reference** |
| --- | --- | --- | --- |
| *Drosophila melanogaster* | “DGRP” founded from 1 outbred population (1500 mated females). | 192/205 | Mackay et al. 2012;<br>Mackay et al. 2018 |
| *Drosophila melanogaster* | “DSPR” founded from 2 outbred populations, created from 15 isolines and recombined for 50 generations, followed by 25 generations of inbreeding | 1600 | King et al. 2012 |
| *Mus musculus* | “RIX” founded from 8 laboratory strains, combined during 3 generations and then inbred during 20 generations. | 69 | Churchill et al. 2004;<br>Srivastava et al. 2017 |
| *Mus musculus* | “BXD ARI” founded from 2 laboratory strains (after 9 to 14 generations of inter-crossing, followed by at least 14 generations of inbreeding | 46 | Peirce et al. 2004 |
| *Caenorhabditis elegans* | Founded from 2 wild-type strains; 15 generations of inbreeding | 73 | van Swinderen et al. 1997 |
| *Caenorhabditis elegans* | From 8 parental lines each initially crossed with males of one different line; 3 to 5 generations of backcrossing followed by 10 generations of selfing | 90 | Doroszuk et al. 2009 |
| *Caenorhabditis elegans* | 12 lines from 2 hybrid populations (6 from each); 6 from 11 generations of selfing and 6 from 22 generations of sib-mating | 12 | Teotónio et al. 2012 |
| *Caenorhabditis elegans* | 359 genotypes from reciprocal crosses between 2 parental lines; 3 generations of backcrossing followed by 10 generations of controlled matings between hybrids | 359 | Andersen et al. 2015 |
| *Caenorhabditis elegans* | From 2 lines; 10 generations of selfing | 120 | Frézal et al. 2018 |
| *Callosobruchus maculatus* | Founded from 215 mated females; 10 generations of inbreeding | ~ 86 (40% of 215) | Bilde et al. 2009 |

Experimental evolution allows following real time adaptation of populations exposed to specific selection pressures (Gibbs 1999, Kawecki et al. 2012). Hence, it allows measuring the process of adaptation itself instead of inferring it based on observed patterns, and to infer causality. Indeed, this method consists in deriving populations from a common ancestral and exposing them to specific controlled environments during several generations, which enables (a) knowledge of the ancestral state of populations, (b) the possibility to define and control the environments that populations are exposed to and (c) replication at the population level (Magalhães and Matos 2012). The explanatory power of experimental evolution can be used to unravel how populations adapt to environmental changes, the presence of antagonists or different population structures (Macke et al. 2011, Kawecki et al. 2012, Rodrigues et al. 2016, Zélé et al. 2018a). Moreover, this method can also be used to measure convergent evolution of different populations to a common environment (e.g. the laboratory; Simões et al. 2008, Fragata et al. 2014).

To ensure that data obtained in the laboratory reflect the range of possible responses found in natural populations of the species under study, the ancestral population should reflect the variability found in the field (MacDonald and Long 2004, Nunes et al. 2008, Faria and Sucena 2017). Moreover, adaptation of non-microbial organisms to rapid environmental changes relies mostly on the standing genetic variation present in a population, rather than on the arrival of new mutations (Hermisson and Pennings 2005, Barret and Schluter 2008, Sousa et al. 2019). Thus, for the establishment of experimental evolution populations, it is crucial to generate and maintain populations with large genetic variability in the laboratory. For this reason, several studies have used laboratory populations founded by a large number of individuals collected in the field and maintained at high numbers in the laboratory (e.g., Mery and Kawecki 2002, Magalhães et al. 2007a, Fricke and Arnqvist 2007, Teotónio et al. 2008, Martins et al. 2013). However, this method falls short on accounting for potential geographical variation in trait values across populations. To ensure this, some studies have produced populations by merging clones, inbred lines (Zbinden et al. 2008, Kover et al. 2009, King et al. 2012), or field populations (Tucic et al. 1995, Fricke and Arnqvist 2007) collected in different locations. This option increases the chance of obtaining a population containing genotypes from different environments, thus potentially representing different subsets of the genetic variability of a species. Yet, this procedure does not preclude the possibility of one (or a set of) genotype(s) from a particular environment invading the whole population. To circumvent this caveat, in sexual organisms, it is desirable to create a population with an equitable representation of the genotypes present in several field populations, which can be achieved by performing controlled crosses between individuals of different populations.

Here, we describe the creation of the above-mentioned biological tools, outbred populations and inbred lines, in two species of haplodiploid spider mites. Additionally, we provide a general description of the experimental evolution protocol we adopted, which can be easily transposed to other systems.

## The system

Spider-mites (Acari: Tetranichidae) are haplodiploid cosmopolitan pests of many agricultural crops of economic interest (Migeon et al. 2010). Given their small size and life-cycle characteristics, these species are easily reared and maintained in high numbers in the laboratory. Therefore, the biology of spider mites is well characterized, as well as their interactions with host plants (Helle and Sabelis 1985, Magalhães et al. 2007b, Kant et al. 2015, Rioja et al. 2017, Bui et al. 2018, Godinho et al. 2018, Paulo et al. 2018). Several experimental evolution studies have been performed with *Tetranychus urticae*, the most studied species. These studies have shown that it rapidly adapts to novel hosts (Fry 1989, Magalhães et al. 2007a, Wybouw et al. 2015, Sousa et al. 2019), to pesticides (Van Leeuwen et al. 2012, Dermauw et al. 2013) and to different population structures (Macke et al. 2011,2012). Additionally, the genome of this species has been sequenced (Grbic et al. 2011), allowing detailed genomic and transcriptomic studies of such adaptation processes (Dermauw et al. 2013, Wybouw et al. 2015, 2019, Jonckheere et al. 2017, Snoeck et al. 2018). *T. urticae* often co-occurs with other spider-mite species (Zélé et al. 2018b). In particular, the congeneric *T. evansi* supresses the defences of several plant species (Sarmento et al. 2011, Alba et al. 2015, Paulo et al. 2018), increasing the performance of con- and heterospecifics on those plants (Sarmento et al. 2011, Godinho et al. 2016). This ability has the potential to strongly affect the ecology and evolution of herbivores and plants that co-occur with this species. Therefore, a study system including both *T. urticae* and *T. evansi* species can effectively tackle several pressing issues in the ecology and evolution of herbivore-plant interactions.

## Collection of spider-mite field populations

To establish highly outbred populations of *T. urticae* and *T. evansi*, several tomato (*Solanum lycopersicum*) fields and greenhouses in Portugal were surveyed for the presence of Tetranychid mites between May and October 2017 (Fig. 1). Each location was sampled during ca. 1 hour. Tomato leaves infested with spider mites were collected and kept in a closed plastic box. If the tomato plants were free of spider mites, neighbouring plants from other species were also screened. All collected spider-mite populations were established in the laboratory by transferring adult females (N= 32 to 463; Table 2) to a rearing cage containing tomato leaves (variety Moneymaker). These populations were maintained under controlled conditions (23.5 ± 2 ºC, 60% RH, 16/8 h L/D) for a few generations (3 to 6), to increase the numbers of spider mites and promote laboratory adaptation. Subsequently, the species was determined for each population by performing a multiplex PCR on a pool of 50-100 spider mites (Zélé et al. 2018d). A total of 27 populations were collected in 24 different locations (Table 2). Sixteen of those were identified as *T. cinnabarinus* (also referred to as the red form of *T. urticae*; Auger et al. 2013), 4 as *T. urticae* (green form) and 7 as *T. evansi*. In 8 locations, there were no spider mites infesting tomato plants but on 4 of those, spider-mite populations were found on neighbouring plants (Table 2).

**Table 2.**
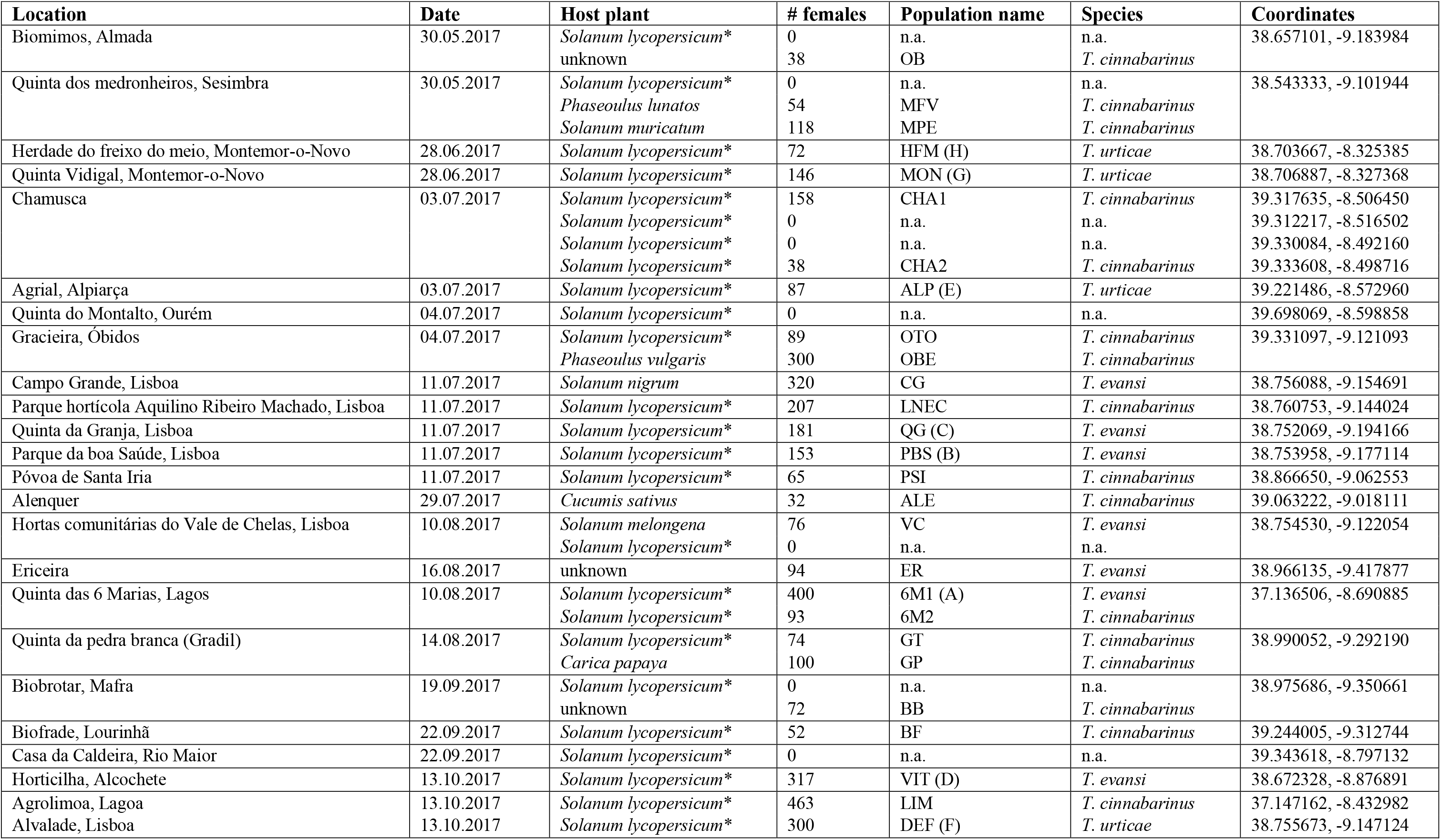
Geographic locations visited to sample spider mite populations. For each location, the table includes the coordinates, the date and the host plants examined, as well as the number of females (# females) and their species when a population was found. n.a. – non-applicable; * tomato.

**Figure 1.**
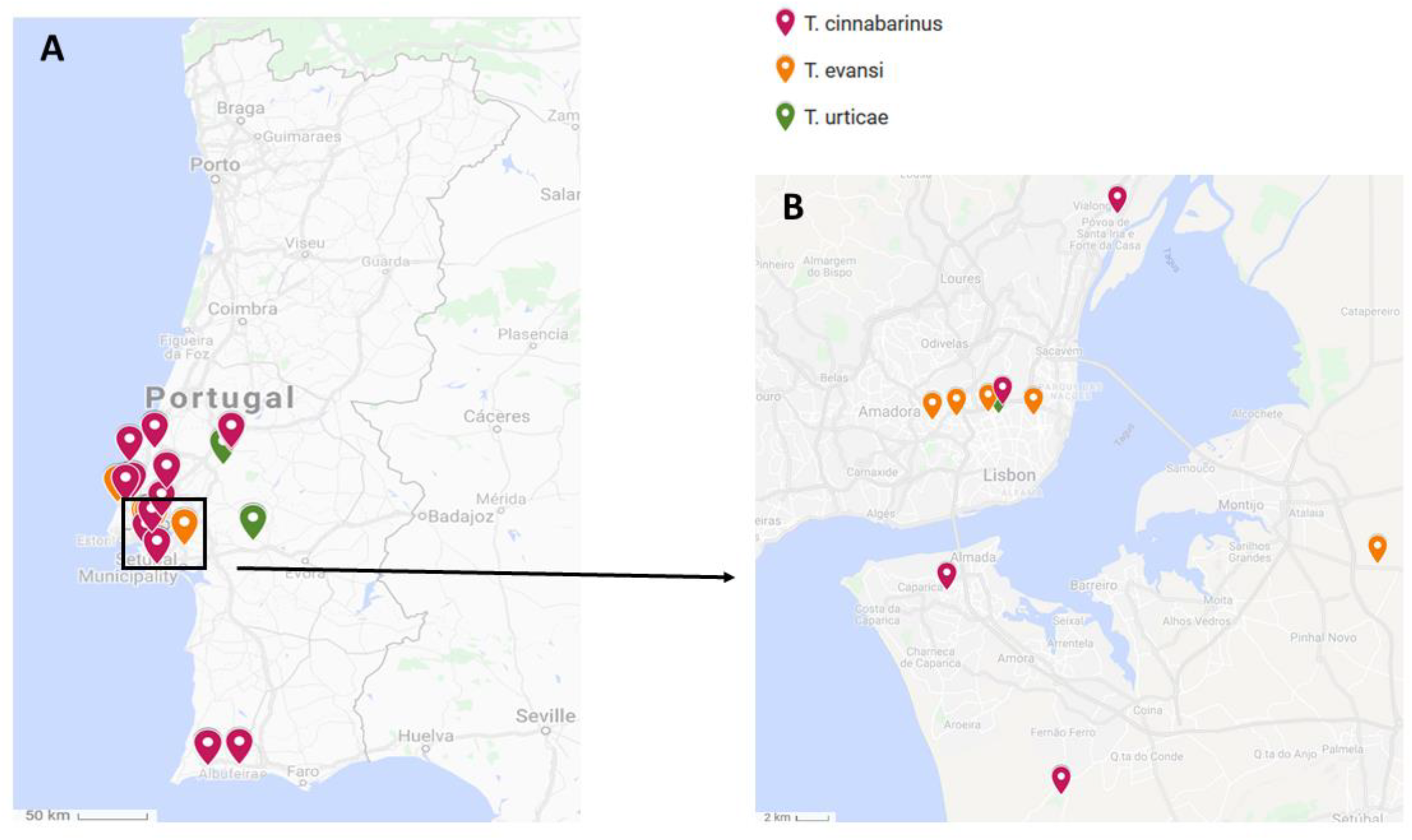
Map of the field sampling of spider mites. A) Total of 24 sites visited in Portugal, B) detailed Lisbon geographic region. This map was adapted from Google Maps (Google n.d. retrieved from https://www.google.com/maps/d/viewer?mid=1v4-9f9Rgmc2HZOfI9TEeoMJB_yS9ZYVu&hl=en&usp=sharing)

## Characterization of populations

We first tested whether the populations collected were resistance to the pesticide etoxazole, as this trait can be used as a genetic marker (Rodrigues et al. 2020). Then, we tested features of the field-collected populations that might lead to reproductive incompatibilities between different populations/genotypes and that could possibly hamper the creation of outbred and inbred populations.

### Resistance to etoxazole

We tested whether most populations collected in the field (7 of *T. evansi*, 4 of *T. urticae* and 3 of *T. cinnabarinus*; Table 3) were resistant to etoxazole. This pesticide interferes with chitin synthesis and deposition, affecting the hatching of embryos and moulting stages (van Leeuwen et al. 2012). To test for resistance to etoxazole, we isolated adult females of each population on individual leaf patches and allowed them to lay eggs for 48 hours. Half of the females of each population (N=31 to 40) were fed on leaf patches placed on water-soaked cotton, while the remaining were fed on leaf patches placed on cotton soaked with etoxazole (500 ppm). After one week, the number of unhatched eggs, alive and dead juveniles was registered. All populations tested showed juvenile survival below 10% when exposed to etoxazole (Table 4), indicating that alleles conferring pesticide resistance, if present, are at very low frequency.

**Table 3.** Infection by endosymbionts and ITS type. Spider mites of each population were tested in a pool (N= 50 to 100) for the presence of endosymbionts (*Wolbachia*, *Cardinium* and *Rickettsia)*. In *T. evansi*, each population was characterized for its ITS type (T1 or T2).

| Species | Population | Symbionts | ITS type |
| --- | --- | --- | --- |
| *T. urticae* | HFM | uninfected | n.a. |
|  | MON | *Wolbachia* |  |
|  | ALP | uninfected |  |
|  | DEF | uninfected |  |
| *T. evansi* | CG | uninfected | T2 |
|  | ER | uninfected | T2 |
|  | PBS | uninfected | T1 |
|  | QG | *Wolbachia* | T1 |
|  | VC | *Wolbachia* | T2 |
|  | VIT | uninfected | T1 |
|  | 6M1 | uninfected | T1 |
| *T. cinnabarinus* | 6M2 | uninfected | n.a. |
|  | LIM | *Wolbachia* |  |
|  | LNEC | *Wolbachia* |  |

n.a. – non-applicable

**Table 4.** Pesticide resistance. Average survival of the offspring of females (N= 31 to 40 per treatment) belonging to different populations of spider mites, exposed to etoxazole or not (control).

| Species | Population | Average juvenile survival |  |
| --- | --- | --- | --- |
|  |  | Control | Etoxazole |
| *T. urticae* | HFM | 0.89 ± 0.04 | 0 ± 0 |
|  | MON | 0.89 ± 0.04 | 0.04 ± 0.03 |
|  | ALP | 0.89 ± 0.04 | 0.09 ± 0.04 |
|  | DEF | 0.92 ± 0.03 | 0.01 ± 0.01 |
| *T. evansi* | CG | 0.94 ± 0.02 | 0.12 ± 0.05 |
|  | ER | 0.90 ± 0.03 | 0.01 ± 0.01 |
|  | PBS | 0.95 ± 0.01 | 0.04 ± 0.03 |
|  | QG | 0.97 ± 0.01 | 0 ± 0 |
|  | VC | 0.88 ± 0.04 | 0.09 ± 0.04 |
|  | VIT | 0.97 ± 0.05 | 0.02 ± 0.01 |
|  | 6M1 | 0.92 ± 0.02 | 0.06 ± 0.03 |
| *T. cinnabarinus* | 6M2 | 0.92 ± 0.06 | 0 ± 0 |
|  | LIM | 0.98 ± 0.01 | 0.08 ± 0.03 |
|  | LNEC | 0.94 ± 0.01 | 0 ± 0 |

### Detection of reproductive incompatibilities among populations

Because the presence of maternally-inherited endosymbionts may hamper the viability of offspring from inter-population crosses (Bordenstein et al. 2001, Telschow et al. 2002, Koukou et al. 2006), we characterized the symbiont community present in most field-collected populations (7 of *T. evansi*, 4 of *T. urticae* and 3 of *T. cinnabarinus*; Table 3). Infection by the most common endosymbionts found in spider mites, namely *Wolbachia*, *Cardinium* and *Rickettsia* (Zélé et al. 2018b), was assessed by multiplex PCR as described in Zélé et al. (2018d). *Wolbachia* was found infecting 6 out of the 14 populations screened, whereas the remaining populations were free of symbionts (Table 3). Subsequently, to avoid incompatibilities among populations due to the presence of endosymbionts, a subset (N>300 females) of each population selected to create the outbred populations (see below) was cured from endosymbiont infection by being continuously exposed to heat shock (33 ºC) for 6 generations (van Opijnen and Breeuwer 1999). Note that due to potential side effects of the heat shock treatment, this procedure was used for all selected populations, independently of whether they were initially infected by symbionts.

To avoid possible reproductive incompatibilities due to genetic differentiation between the green and the red forms of *T. urticae* (e.g. Deboer 1982, Gotoh and Tokioka 1996, Sugasawa et al. 2002), we only used populations of the green form and discarded most populations of the red form (or *T. cinnabarinus*) after genetic identification. Additionally, in *T. evansi*, two highly incompatible major clades, I and II, have been identified based on the cytochrome oxidase complex I (COI) haplotypes and the internal transcribed spacer region (ITS; Boubou et al. 2011; Knegt et al. 2017). To avoid such incompatibility, we sequenced (Stabvida, Caparica, Portugal) the ITS of *T. evansi* populations (Table 3) and used only the populations with ITS type T1, which corresponds to clade I, to create the outbred population.

## Creation of outbred populations in haplodiploids

To create outbred populations with high levels of standing genetic variation, we used 4 symbiont-free populations of each species belonging to the same haplotype (the green form for *T. urticae* and Clade I for *T. evansi*). These populations were 6M1, PBS, QG and VIT for *T. evansi* and ALP, DEF, MON and HFM for *T. urticae* (Table 2; hereafter labelled A to D and E to H, respectively). These populations were then merged by performing inter-population crosses in a controlled fashion, to avoid over-representation of genotypes from a given population (Figure 2). To this aim, 200 females from population A were individually placed on leaf patches (Ø18 mm) and crossed with 200 males from population B and vice versa. Additionally, 200 females from population C were individually crossed with 200 males from population D and vice versa. In this way, we obtained a hybrid F1 (AB, BA; CD and DC). However, because spider mites are haplodiploid with an arrhenotokous genetic system (Helle and Sabelis 1985), only the F_1_ female offspring resulting from these crosses are hybrids, as sons stem from unfertilized eggs. To form hybrid males, virgin hybrid F_1_ females were collected during their last moult, allowed to emerge as adult female and to lay unfertilized eggs for 48h. Subsequently, their offspring (haploid hybrid males) was allowed to develop until adulthood. To synchronize the generations at which hybrid females and males were produced, a new generation of F_1_ hybrid females (again AB, BA, CD and DC) was obtained by repeating the previous set of matings one generation later. These hybrid females were then crossed with hybrid males to produce a fully-hybrid F_2_ (AB and BA hybrid females were crossed with CD or DC hybrid males and vice versa). Again, because males stem from unfertilized eggs, only the female offspring resulting from these crosses was a fully-hybrid combination of the 4 populations (e.g. ABCD). These fully-hybrid females were also isolated as virgin and their sons allowed to develop until adulthood. To synchronize the production of fully-hybrid adult males and females, another cross of AB and BA females and CD or DC males (and vice versa) was performed simultaneously (Fig. 2a). Finally, individuals of both genders of each of the 8 fully-hybrid combinations performed (ABCD, ABDC, BACD, BADC, CDAB, CDBA, DCAB and DCBA) were mixed to form the outbred population (Fig 2a).

**Figure 2.**
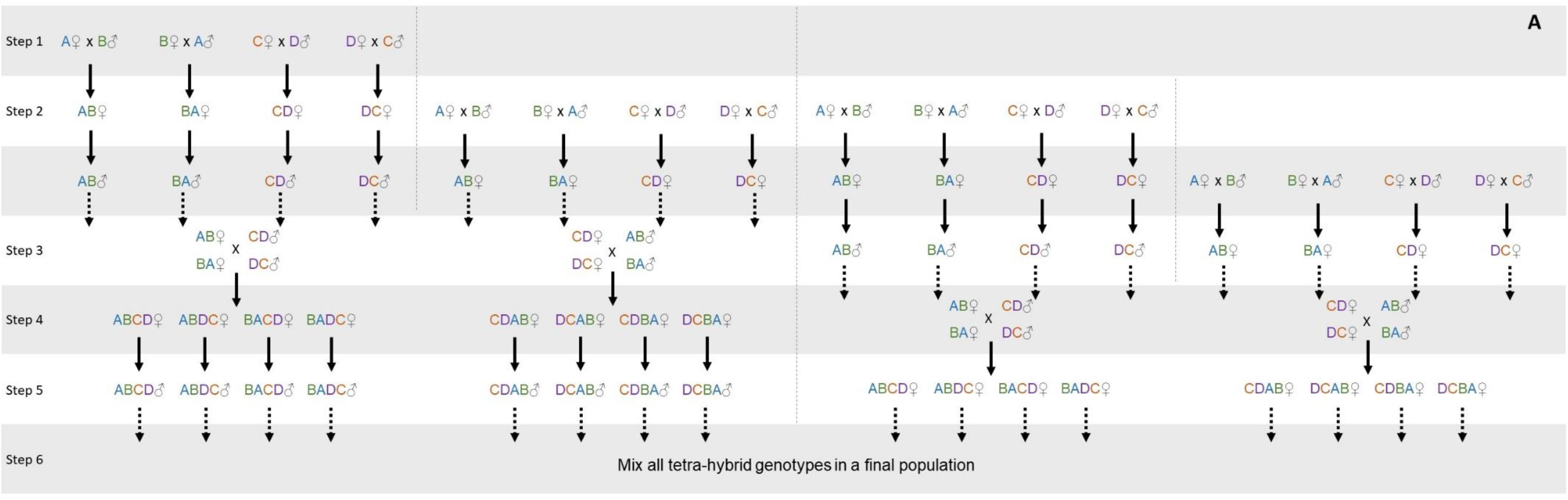
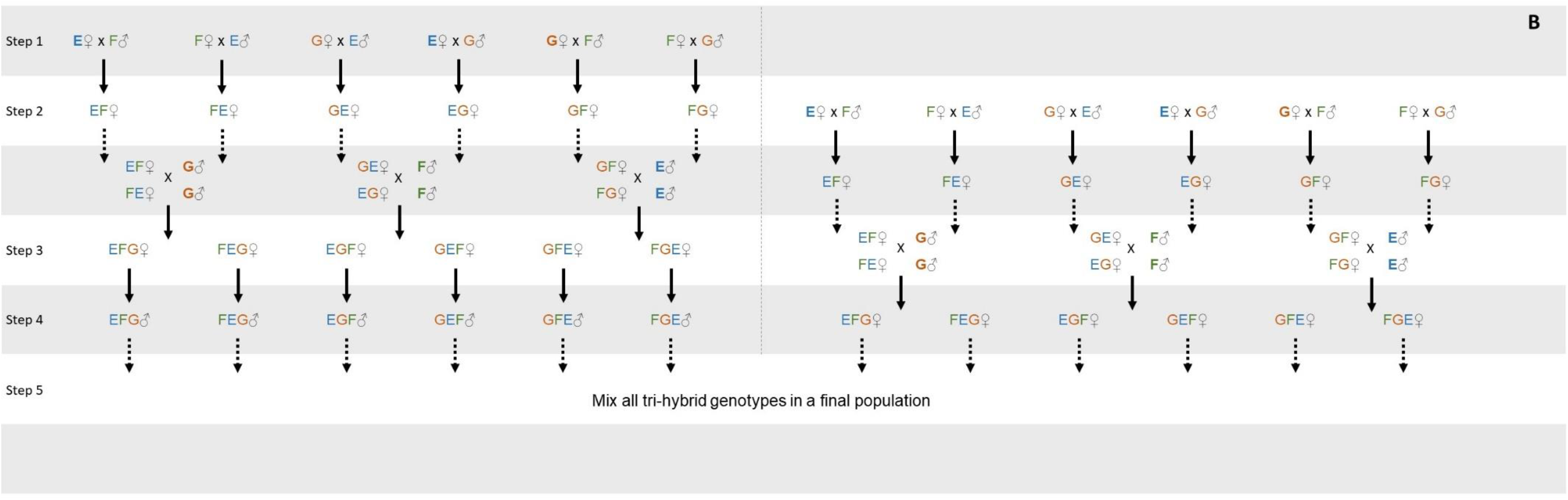
Creation of outbred populations on haplodiploids. An outbred population of haplodiploid spider mites was created by performing controlled crosses between **A** four or **B** three populations collected in different locations. Letters represent the population from which the genotypes derive from. Each step represents the production of offspring to use in the crosses for the following generation: females are obtained from crosses between different genotypes and males are obtained from virgin females of a given genotype. Bold arrows represent the development of the offspring forming the next generation, dashed arrows represent the use of hybrids for the subsequent crosses.

This procedure was done to ensure an equal genetic representation (nuclear and mitochondrial) of each field population in the resulting outbred population. Because some crosses did not produce viable offspring of at least one sex, we opted to found the outbred population with an equal number of individuals from each different combination (ex. ABCD), corresponding to the minimum number of genotypes obtained in those combinations. This means that the *T. evansi* outbred was founded with 72 females and 72 males from each of the 8 combinations performed (a total of 576 different genotypes from 4 locations).

During the creation of the *T. urticae* outbred population, hybrid breakdown was detected between the population HFM and the three others (i.e. ca. 75% of F_2_ offspring were inviable). The protocol was then adapted to merge 3 populations instead of 4 (Fig. 2b), and the outbred population was founded with 51 females from each of 6 different combinations, corresponding to a total of 306 females. Because the total number of males was low (N=197), we opted to use them all, even though the number across genotypes was not even.

## Creation of inbred lines in haplodiploids via full sib mating

We created 59 inbred lines of *T. evansi* through 15 generations of sib-mating. Below we describe the methods used to create these lines, as well as an estimation of their inbreeding coefficient.

Inbred lines were initiated by isolating a mated female randomly sampled from the outbred population. Given full first-male sperm precedence in spider mites, all descendants of a female stem from the same father (Rodrigues et al. 2020), which reduces genetic variance in the offspring as compared to species with mixed paternity. In this case, and given that spider mites are haplodiploid, a maximum of three different alleles (e.g. x, y and z) can be initially sampled at each locus, independently of the number of alleles available for that locus in a population. Hence, 4 different types of mated females can be sampled, which correspond to 4 possible types of crosses: (A) a heterozygous female mated with a male that does not share any allele with her (e.g. [xy]*[z]); (B) a heterozygous female mated with a male that shares one of her alleles (e.g. [xy]*[x]); (C) a homozygous female mated with a male that does not share an allele with her (e.g. [xx]*[y]) or (D) a homozygous female mated with a male that has the same allele as her for that locus (e.g. [xx]*[x]).

Because we do not have access to the genotype of the mated female that initiated each line, we assume the most heterozygotic situation, i.e. that we collected a [xy] female mated with a

[z] male. By doing so, we conservatively underestimate the coefficient of inbreeding. This cross (type A) will produce ½ [xz] + ½ [yz] females and ½ [x] + ½ [y] males in the F_1_. If these daughters and sons mate randomly among themselves, crosses among sibs will occur with the following probability: ¼ [xz]*[x] (type B) + ¼ [xz]*[y] (type A) + ¼ [yz]*[x] (type A) + ¼ [yz]*[y] (type B). Thus, at each time step, type A crosses will result in sons and daughters that, if they mate randomly among themselves, will produce ½ type A crosses and ½ type B crosses. Following the same reasoning, type B crosses, e.g. [xy]*[x], will produce ½ [xx] + ½ [xy] females and ½ [x] + ½ [y] males. If these daughters and sons mate randomly, crosses among siblings will occur with the following probability: ¼ [xx]*[x] (type D) + ¼ [xx]*[y] (type C) + ¼ [xy]*[x] (type B) + ¼ [xy]*[y] (type B). Thus, type B crosses will result in sons and daughters that, if they mate randomly among themselves, will produce ½ type B crosses, ¼ type C crosses and ¼ type D crosses. From a type C cross, e.g. [xx]*[y], only [xy] females and [x] males will be produced and, thus, only [xy]*[x] crosses (type B) will occur. Finally, type D crosses, e.g. [xx]*[x], will only produce [xx] females and [x] males, and, thus only type D crosses will occur.

Therefore, the frequencies of each type of cross, at each generation (t+1), are given by the following equations:

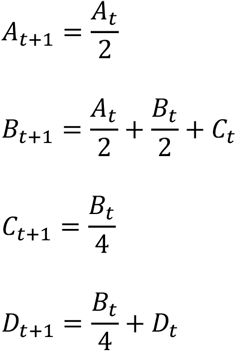

The coefficient of inbreeding (*f*_*t*_), which corresponds to the probability that two alleles at one locus are identical by descent (Wright 1921), can subsequently be obtained using the following equation:

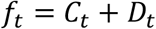

Alternatively, for full sib-mating in haplodiploids, this coefficient can also be obtained directly as:

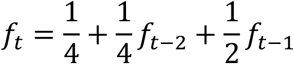

where the first two terms correspond to the probability of both alleles coming from the grandmother, being the alleles equal (first term), thus, f=1, or different (second term), thus, f is equal to that of the grandmother f_(t-2)_ and the third term corresponds to the probability of one allele coming from the grandmother and the other from the grandfather (i.e. f is the same as that of the mother f_(t-1)_).

Both methods yield the same result, and assuming that generation 0 starts with a [xy] female mated with a [z] male (i.e. *A*_0_ = 1 with the first method and *f*_1_ = *f*_2_ = 0 with the second method), we obtain a coefficient of inbreeding of 95.1% after 15 generations (Fig. 3). However, the first method also allows estimating the probability of having a fully inbred line, which is given by the frequency of individuals stemming from fully homozygous crosses (*D*_*t*_). Again, assuming the most heterozygotic scenario, we obtain a probability of having a fully inbred line of 93.6% after 15 generations (Fig. 3).

**Figure 3.**
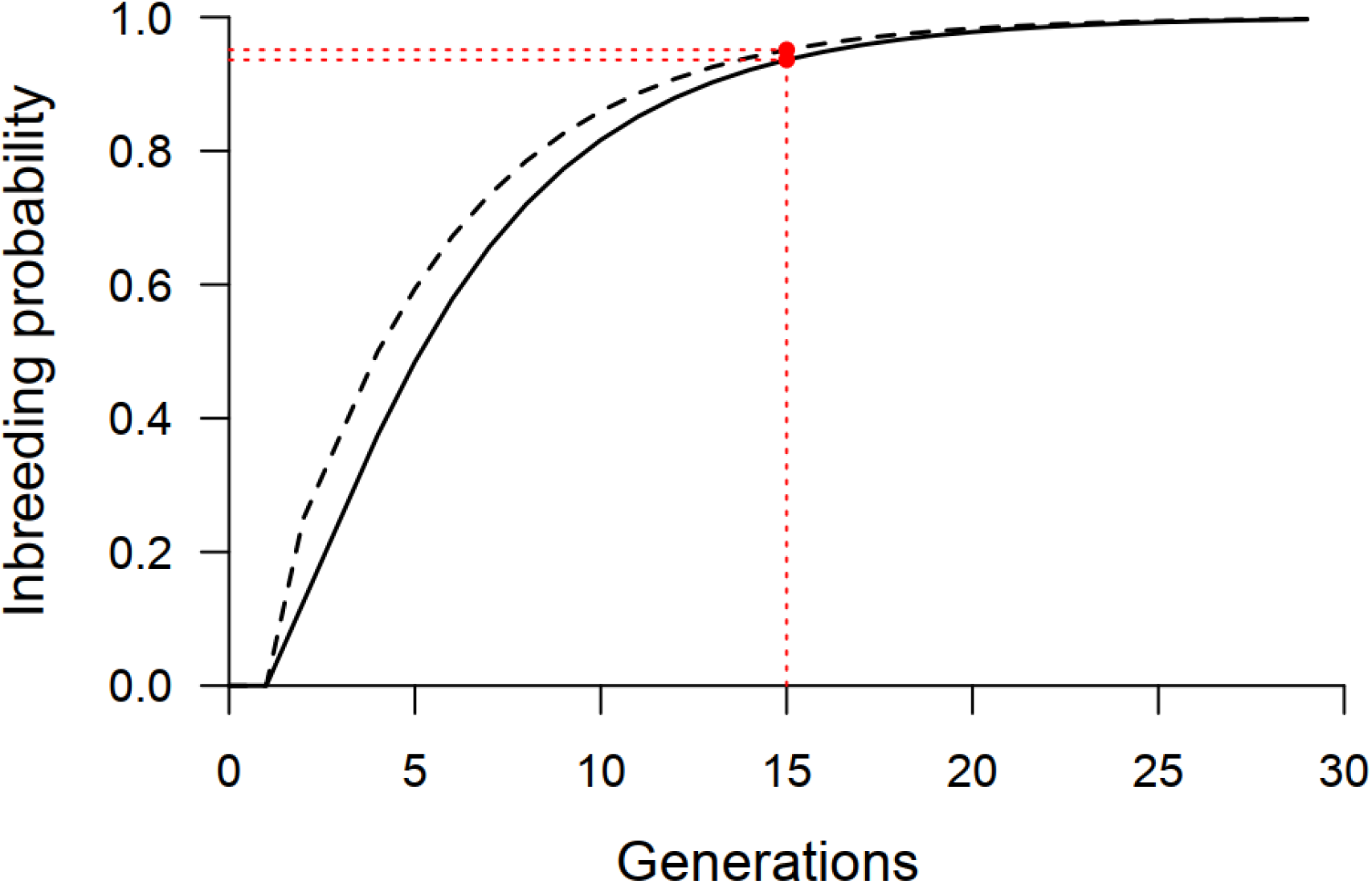
Inbreeding estimates from full sib-mating crosses in haplodiploids. Coefficient of inbreeding (f_t_; dashed line) and probability of having a fully inbred line (D_t_; full line) for each discrete generation of full sib-mating, starting from the most heterozygotic combination at one locus (e.g. a [xy] female mated with a [z] male).

To create inbred female lines from the *T. evansi* outbred population, we randomly sampled 450 mated females and installed them individually on tomato-leaf patches (Ø18 mm), where they laid eggs for 48h. The offspring of each female was then allowed to develop until adulthood (10 to 12 days) and to mate on that patch (i.e. sib-matings). After 14 days, 3 mated females from each patch were isolated on 3 new patches and the same procedure was repeated. On the following generation, 3 sib-mated females from one of the three patches only, were isolated on 3 new patches and allowed to oviposit for 48h. The entire procedure was then repeated for 15 discrete generations. Having 3 replicates per line decreases the chances that lines are lost at each generation. However, despite this, many lines were lost due to the death of the female, null fecundity, no egg hatching, or no female or male offspring produced by a given female (Fig. 4). After 15 generations of sib-mating, each of the remaining inbred lines (N = 59) was transferred to individual patches of tomato plants kept on water-soaked cotton in petri dishes and maintained in small numbers thereafter.

**Figure 4.**
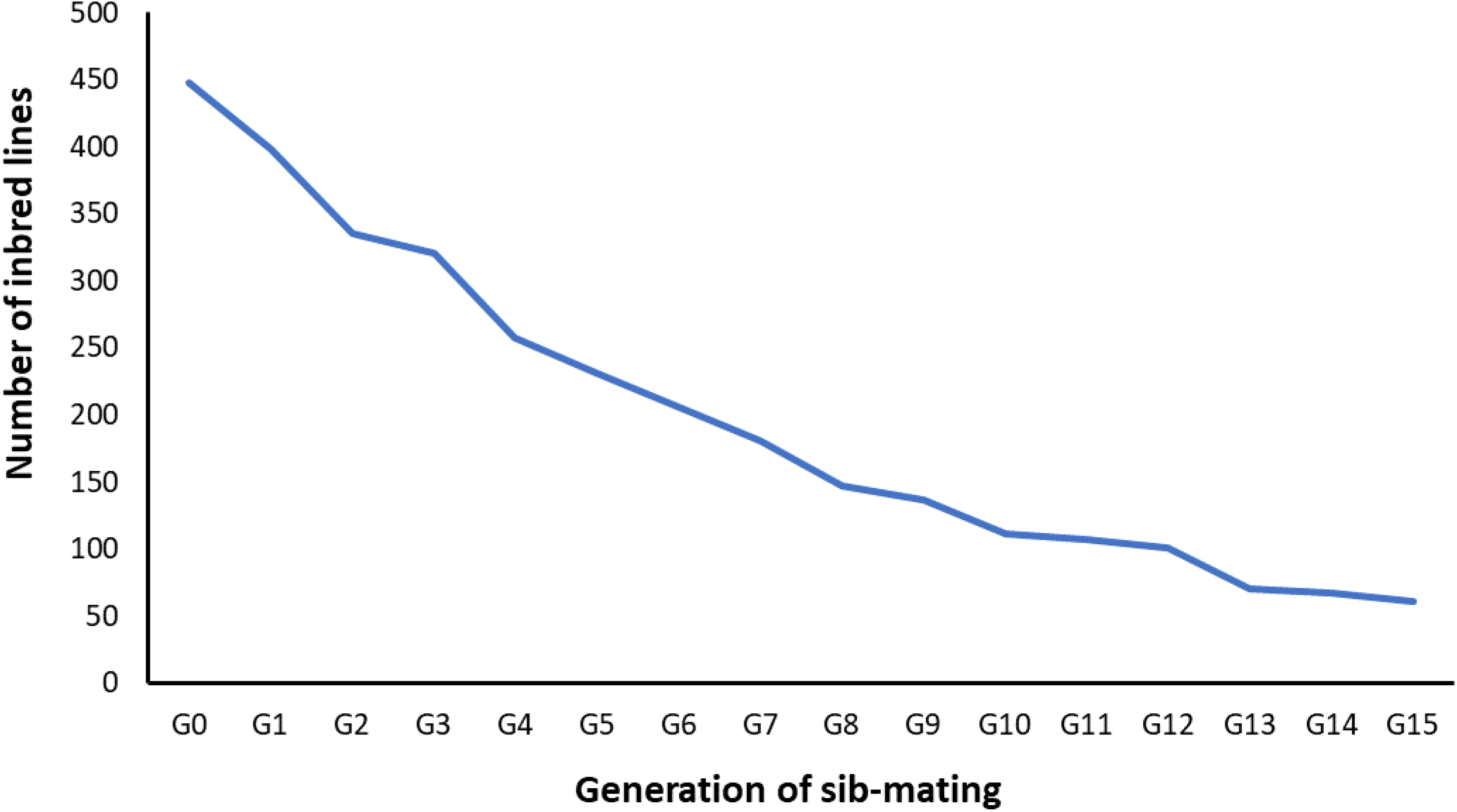
Creation of inbred lines in haplodiploids. Number of inbred lines of *T. evansi* across 15 generations of sib-mating.

## Experimental evolution protocol

The experimental evolution protocol was initiated by transferring 220 randomly collected females from the outbred population (to ensure >200 living females founding each experimental population) to a box corresponding to a given selection regime. This procedure was repeated 5 times per selection regime, as the replicate unit in experimental evolution studies is the population (Kawecki et al. 2012). All populations were maintained under the same environmental conditions except for variables that corresponded to each selection regime (e.g., type of host plant, presence/absence of competitors). Each generation, 220 randomly selected mated females were transferred to a new box with the same characteristics. The remaining individuals were maintained in the original box until the next transfer, creating a back-up population (t-1) for each replicate of each selective regime. Thus, if the 220 females could not be found in a given box at the time of transfer, the remaining number was transferred from the back-up population. If the sum of females found in the experimental population and its respective t-1 back-up was not sufficient, the remaining number of mites was transferred from the base outbred population to reach 220. This procedure allowed maintaining the same population size for each replicate. However, because every generation each experimental population could be exposed to a given number of migrants from the t-1 back-up population and/or the base population, the effective number of generations of selection might differ. This number was estimated using the following equation:

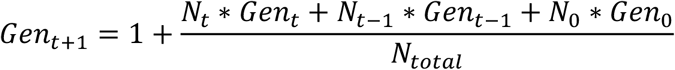

Where Gen_t+1_ corresponds to the effective number of generations of selection underwent by the individuals at the next generation. Gen_t_, Gen_t-1_ and Gen_0_ correspond, respectively, to the effective number of generations of selection underwent by the current generation, the previous generation and the base populations (i.e. not adapted to the new environment, so Gen_0_=0). N_t_, N_t-1_ and N_0_ correspond to the number of individuals transferred from the current generation of selection, the back-up t-1 box and the base population, respectively, and N_total_ to the total number of adult females transferred. We provide an example of this formula by applying it to our system in Fig. 5.

**Figure 5.**
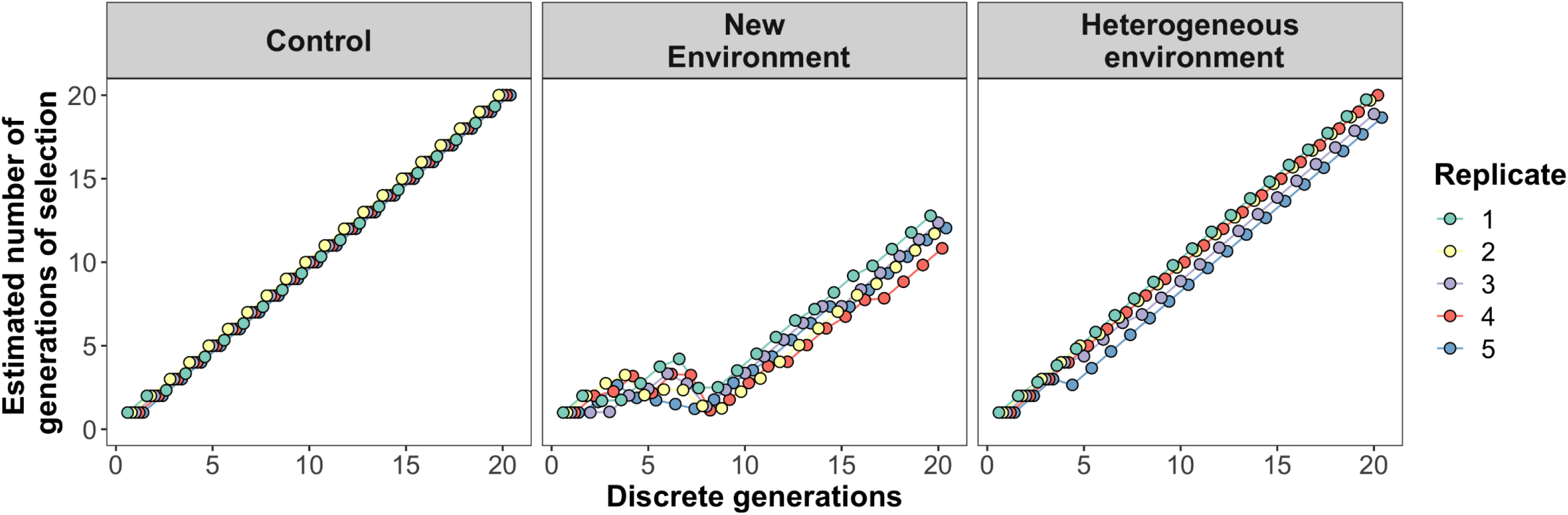
Estimated number of effective generations of selection. Populations were exposed to an environment similar to that of the ancestral population (control), to a new environment, or to a mixture of both (heterogeneous environment). Because individuals from the t-1 and base populations were often added in the selection regime corresponding to a new environment, the estimated number of generations decreases considerably relative to the other selection regimes. However, this procedure allowed populations to overcome the initial reduction in population size and to subsequently adapt to the selection regime imposed.

## Discussion

We describe the creation of tools that maximize the maintenance of standing genetic variation in laboratory populations of haplodiploids and thus allow characterizing responses to selection pressures. As a case study, we present the creation of outbred populations for two spider mite species, *T. urticae* and *T. evansi*, by performing controlled crosses between recently collected field populations. In addition, we report a procedure to calculate the inbreeding coefficient, but also the probability of having a full inbreed line when performing full sib-mating in haplodiploids and applied it for the creation of inbred lines of *T. evansi*. Finally, we provide an outline of the experimental evolution protocol with a methodology that allows maintaining constant densities across generations of selection.

Undoubtedly the method we present here is very time and work consuming. However, we believe that the advantages of creating such powerful tools, as we describe here, compensate the effort in the long run. Additionally, the time spent with the creation of outbred populations gives the populations time to adapt to the laboratory conditions, a requirement before testing adaptive responses to other environments (Matos et al. 2002, Simões et al. 2008, Fragata et al. 2014). Moreover, while developing these tools, the populations collected may be thoroughly characterized, providing a preview of the variability expected in the derived outbred population and inbred lines.

As populations were collected from nearby locations, the resulting populations do not encompass large areas of potential geographical variation, unlike for example the populations used to create the DSPR panel, where the 15 founder genotypes have distant geographic origins (King et al. 2012). This may limit the standing genetic variation available, because of similar environmental conditions, and/or migration among populations. However, this is not very likely in the case of spider mites, as (a) experimental evolution studies performed with populations from a single location have repeatedly shown responses to selection (reviewed in Sousa et al 2019), and (b) populations of spider mites show high genetic differentiation even within small geographical scales (e.g. Bailly et al. 2004, Carbonelle et al. 2007). Additionally, by founding the outbred populations with more than 500 individuals, the chances of obtaining large amounts of standing genetic variation are high.

Performing controlled crosses among field populations maximizes the chances of obtaining a highly outbred laboratory population. Indeed, this method allows controlling for assortative mating and for differences in fitness and/or mating competitive ability between genotypes. Therefore, it ensures an equal genetic representation of the nuclear and mitochondrial genomes of each population by equalizing the number of genotypes from each field population that will be incorporated in the outbred population. Finally, it allows the detection of reproductive incompatibilities between populations/genotypes, and the consequent exclusion of inviable crosses. Indeed, throughout the process of creation of these outbred populations, several crosses resulted in infertile females, or inviable offspring, indicating possible incompatibilities. Specifically, we detected hybrid breakdown (i.e. postzygotic reproductive isolation where F_2_ hybrids are inviable or sterile) between one population of *T. urticae* (HFM) and the others. None of these potential causes of reduction of the genetic variability in the final population would have been detected and prevented if the outbred population had been founded by mixing several individuals of different populations without controlling for the outcome of such crosses at the individual level.

Using outbred populations increases the chance that the responses observed are representative of the study species, which is a common shortcoming of laboratory studies. For example, Vala et al. (2004) found that *T. urticae* females that are not infected with *Wolbachia* prefer uninfected over infected males, thereby potentially reducing the costs of incompatible mattings. However, this result was based on a single line, whereas a later study using an outbred population stemming from several field populations does not recapitulate this result (Rodrigues et al. biorxiv). Another example concerns the interaction between *T. urticae* and tomato plant defences. Although this herbivore generally induces plant defences, some field-collected lines were shown to supress them instead (Kant et al. 2008). Therefore, capturing and maintaining natural variation in laboratory studies is highly relevant for understanding the ecology and evolution of the interaction between study organisms, such as spider mites, and many environmental factors, such as symbionts or plants.

In our study system, several experimental evolution studies have been performed using *T. urticae* (e.g. van Leeuwen et al 2012, Masier and Bonte 2019, Sousa et al. 2019). However, these studies have been performed with populations or strains collected from a single location, and in some cases from a small number of individuals (reviewed in Sousa et al. 2019). Therefore, the responses obtained may be idiosyncratic of the genetic background used. Thus, providing the community with highly outbred laboratory populations may be very useful to test the generality of the responses reported, and to perform future studies on other topics with a larger representation of the genetic variation of the species. Moreover, as genetic variance is the raw material for selection to act upon, having a highly outbred population to initiate experimental evolution will increase the chances of observing fast responses to selection.

Clearly, performing laboratory studies with outbred populations adds to their robustness. However, to characterize the potential different responses found in such outbred populations due to their genetic variance, it is necessary to fix this variance along a panel of inbred lines, such that each genotype can be studied independently. In particular, studying these lines will allow a clear understanding of the phenotypic and genotypic variability for traits that may be relevant in many different contexts. Importantly, inbred lines can also be used to assess genetic correlations and trade-offs between different traits (e.g. Travers et al. 2015, Howick and Lazzaro 2017, Wang et al. 2017, Lafuente et al. 2018, Everman et al. 2019), including those measured in different environments (Howick and Lazzaro 2014, Unckless et al. 2015, Orsted et al. 2018). Indeed, because all individuals of a given inbred line represent roughly the same genotype, responses of each genotype can be measured in different contexts. Additionally, such inbred lines can be used as a fixed genetic background against which the response of another population is studied. This may be particularly useful in the context of the evolution of biological interactions. For example, unravelling the evolution of sexual conflicts can be done by exposing individuals of the evolving sex to inbred lines of the non-evolving sex (e.g. Macke et al. 2014). Also, the magnitude of GxG (genotype by genotype) in host parasite interactions has been addressed by exposing different lines of hosts and/or parasites to each other (Lambrechts, Fellous and Koella 2006, de Roode and Altizer 2010, Carpenter et al. 2012, Hudson et al. 2016). Finally, there is a recently increasing interest on how inter-individual variation affects several ecological characteristics, such as species persistence and coexistence (Lichstein et al. 2007, Agashe 2009, Lankau 2009, Bolnick et al. 2011, Forsman and Wennersten 2016, Hart et al. 2016). Within this context, the creation of inbred lines may also be a useful tool.

Here we also provide an outline of an experimental evolution protocol. Of particular interest is the fact that we have kept populations from the previous generation of each replicate of the different selection regimes. Individuals from these populations can then be used to populate the current generation of experimental evolution when necessary. In this way, it is possible to avoid losing replicates, as it commonly occurs in experimental evolution studies. Moreover, environments that impose a strong selection pressure are, by definition, harsh. Hence, there is a high probability that the populations adapting to those conditions crash in a few generations. Thus, here we propose a method that allows using populations from the previous generation of selection to ensure that the total population size remains constant across generations, thereby allowing populations to overcome the initial reduction in population size. Such populations can thus be rescued and subsequently adapt to the selection regime imposed (Fig. 5). In this case, it is important to, calculate the effective number of generations of selection that populations have undergone, such as to correctly compare responses among selection regimes. Note that these t-1 populations can be kept under relaxed conditions, thereby minimizing the workload necessary to maintain them.

Finally, having the same background in the outbred population and the inbred lines allows comparing results stemming from both types of populations when tackling a common question. The biological resources described in this study can also be shared with several collaborating laboratories and combined with the increasingly fast advances on the genetic and genomic resources available. This will allow consistent and comparable studies that unquestionably will provide great advances in many different frameworks.

## Acknowledgements

We would like to thank Margarida Matos and Élio Sucena for important inputs on the design of the crosses for the creation of the outbred populations, Lucie Sousa and Inês Santos for growing the plants and keeping the field-collected populations and all the members of the MITE2 group for stimulating discussions.

This study was funded by an ERC Consolidator Grant (COMPCON, GA 725419) to SM and by an FCT Ph.D. scholarship PD/BD/114010/2015 to DPG.

## Author’s contribution

DPG and SM designed the study with help from LRR and FZ. DPG and LRR collected the spider mite populations. The creation of the outbred and inbred populations was performed by DPG, MAC, MCM, JTP and CE. FZ and IF developed the formula to calculate the coefficient of inbreeding and the effective number of generations of selection. The manuscript was written by DPG and SM with considerable contributions from all authors.

